# Oropouche orthobunyavirus infection is mediated by the cellular host factor Lrp1

**DOI:** 10.1101/2022.02.26.482111

**Authors:** Madeline M. Schwarz, David A. Price, Safder S. Ganaie, Annie Feng, Nawneet Mishra, Ryan M. Hoehl, Farheen Fatma, Sarah H. Stubbs, Sean P.J. Whelan, Xiaoxia Cui, Takeshi Egawa, Daisy W. Leung, Gaya K. Amarasinghe, Amy L. Hartman

## Abstract

Oropouche orthobunyavirus (OROV; *Peribunyaviridae*) is a mosquito-transmitted virus that causes widespread cases of human febrile illness in South America, with occasional progression to neurologic effects. Host entry factors mediating cellular entry of OROV are undefined. Here, we show that OROV requires the host protein low density lipoprotein-related protein 1 (Lrp1) for cellular infection. Cells from evolutionarily distinct species lacking Lrp1 were less permissive to OROV infection than cells with Lrp1. Treatment of cells with either the high-affinity Lrp1 ligand receptor-associated protein (RAP) or recombinant ectodomain truncations of Lrp1 significantly reduced OROV infection. Thus, Lrp1 is an important host factor required for infection of cells by OROV. Taken together with our results showing that Lrp1 is a newly identified receptor for Rift Valley fever virus (RVFV), Lrp1 may play a broader role in cellular infection by bunyaviruses than previously appreciated.

## Introduction

Bunyaviruses are a large group of related viruses with single-stranded, segmented, negative or ambisense RNA genomes (Abudurexiti et al., 2019). Within the order *Bunyavirales*, the *Peribunyaviridae* family contains viruses that infect humans and animals with confirmed or potential zoonotic transmission (Knipe and Howley, 2013; McMullan et al., 2012). Oropouche virus (OROV; Orthobunyavirus genus; Simbu serogroup) is found primarily in the South American regions of Brazil, Trinidad, Peru, Panama, and Tobago (Sakkas et al., 2018). OROV has caused more than 30 epidemics resulting in over 500,000 total cases of human febrile illness, making it the second most common arboviral disease in Brazil behind Dengue fever (Mouraao et al., 2009; Pinheiro et al., 1981; Sakkas *et al*., 2018). The true case number is likely higher as clinical testing for OROV is lacking and patients are often misdiagnosed as having Chikungunya or Dengue fevers (Sakkas *et al*., 2018). Arthropod vectors for OROV include *Culicoides* midges and *Culex* mosquitoes. In people, OROV causes a debilitating febrile illness that manifests as fever, intense headache, myalgia, joint pain, retro-orbital pain, and photophobia which can further develop into encephalitis or meningitis (Mouraao *et al*., 2009; Pinheiro et al., 1982; Pinheiro *et al*., 1981). Systemic infection manifests as rash, nausea, vomiting, and diarrhea. Viremia and leukopenia are common features (Pinheiro *et al*., 1981), and virus can be detected in the cerebral spinal fluid (Bastos Mde et al., 2012; Chiang et al., 2021). In mice, the virus replicates in the liver and spleen after either subcutaneous or intracranial infection (Proenca-Modena et al., 2016).

Due to the broad cellular tropism and ability to infect a variety of species, bunyaviruses are thought to use multiple receptors or attachment factors for entry and/or a protein that is broadly expressed across different tissues and widely conserved across species. Recently, using a CRISPR/Cas9 screen, we reported that the conserved host protein low-density lipoprotein receptor-related protein-1 (Lrp1) mediates cellular infection with Rift Valley fever virus (RVFV), a phlebovirus within the *Bunyavirales* order (Ganaie et al., 2021). Lrp1 (also known as alpha-2-macroglobulin receptor or CD91) is a highly-conserved multi-functional member of the LDL receptor family of transmembrane surface proteins. Lrp1 is important for ligand endocytosis, cell signaling, lipoprotein metabolism, blood-brain barrier maintenance, angiogenesis, and entry of viruses and bacterial toxins (Gonias and Campana, 2014; Heissig et al., 2020; Potere et al., 2019) (Bres and Faissner, 2019). Homozygous deletion of Lrp1 is embryonically lethal in mice (Herz et al., 1992), further supporting the critical nature of Lrp1 in homeostatic functions.

The M segment of *Bunyavirales* encodes the surface glycoproteins Gn and Gc which form heterodimers and multimerize on the surface of the virion. Few studies have been conducted on the binding and entry mechanisms of OROV Gn/Gc. Given the conserved nature of Lrp1 and its broad expression in different tissues, we asked if OROV, a bunyavirus distantly related to RVFV, also requires Lrp1 to infect host cells. Here, we used Lrp1 knockout (KO) cell lines to show that OROV infection is decreased compared to parental cells expressing Lrp1. Pre-treatment of cells with varying concentrations of the Lrp1-binding protein RAP or Lrp1 ectodomains significantly reduced OROV infection. Zika virus (ZIKV), an arbovirus outside the *Bunyavirales* order, was unaffected by the loss of Lrp1 or by treatment with Lrp1-binding RAP protein. These findings highlight the importance of Lrp1 in OROV orthobunyavirus infection.

## Results

### OROV infection is reduced in Lrp1 knockout (KO) cell lines

OROV strain BeAn19991 (Acrani et al., 2015) was grown in mouse microglial BV2 cells at a multiplicity of infection (MOI) of 0.1 and 0.01 along with RVFV strain ZH501 and ZIKV strain PRVABC59 for comparison (**Fig. S1A**). While ZIKV did not replicate well in BV2 cells, OROV and RVFV reached 10^6^ PFU/mL by 24 hours post-infection (hpi) at MOI 0.1, and these parameters were used for the remaining cellular infection studies. Clonal BV2 knockout (KO) cell lines that are deleted for either Lrp1 or the Lrp1 chaperone protein RAP express significantly reduced levels of Lrp1, and this was visualized and quantified using immunofluorescence and western blot (**Fig. S1B-D**) (Ganaie *et al*., 2021). Infection of both Lrp1 and RAP clonal KO cell lines with OROV resulted in significantly less infectious virus produced by 24 hpi when compared to infection of BV2 wt cells (**Fig. 1A**), with reductions of 2-3 log in titer for both OROV and RVFV as a comparator. Thus, OROV requires Lrp1 or related proteins for efficient cellular infection and production of infectious virus from BV2 cells.

**Figure 1.**
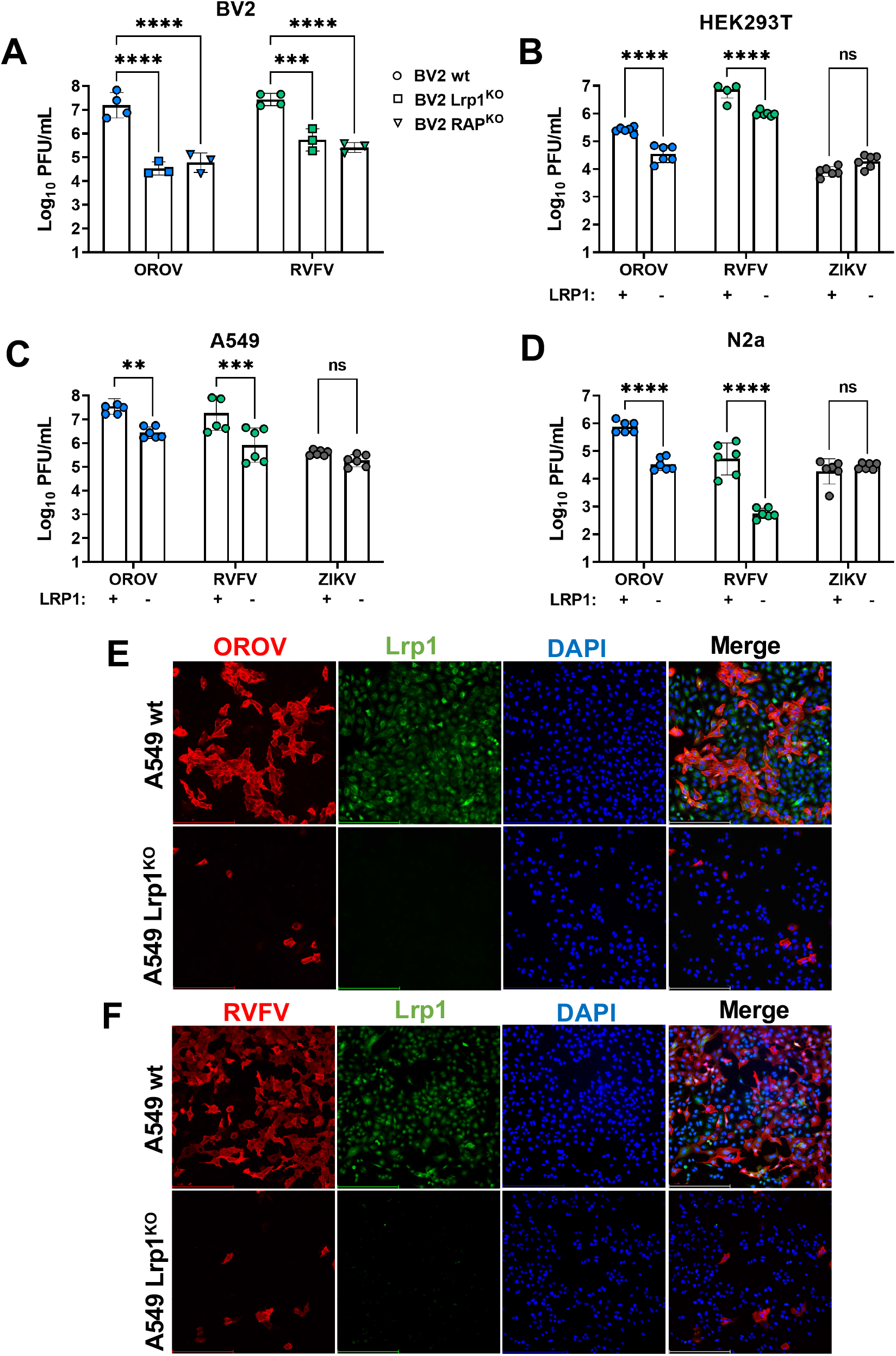
OROV and RVFV show reduced infection in multiple cell lines that are KO for Lrp1. (A) Infection of Lrp1KO and RAPKO cell lines described in Ganaie et al. 2021 with RVFV and OROV at MOI 0.1. Infection of wt (+) and Lrp1 KO (-) versions of (B) HEK293T, (C) A549, and (D) N2a cells with OROV, RVFV, and ZIKV at MOI 0.1. OROV and RVFV samples were harvested at 24 hpi and infectious virus was measured by viral plaque assay. ZIKV samples were harvested at 48 hpi and vRNA was evaluated by qRT-PCR. Fluorescent microscopy (20X) of (E) OROV and (F) RVFV infection of A549 wt and Lrp1 KO cells at MOI 0.1 at 24 hpi. Scale represents 250μm. Statistical significance was determined using an unpaired t-test on log-transformed data. Experiments were repeated three times.**, p<0.01; ***, p<0.001; ****, p<0.0001.

Additional clonal Lrp1 KO cell lines were established in human HEK293T, A549, and murine N2a cell lines, with the loss of Lrp1 verified by western blot **(Fig. S1D).** As with BV2 cells, Lrp1 KO resulted in significantly reduced OROV infection across all cell lines. While similar reductions were seen with RVFV in Lrp1 KO cells, no significant difference in virus infection or production was observed in cells infected with ZIKV, a flavivirus used as a control (**Fig. 1B-D**). The titers for both OROV and RVFV infection of KO cell lines was 10-100 fold lower than wt cell lines. By immunofluorescence, Lrp1 was detectable in wt parental A549 cells but was absent from Lrp1 KO (**Fig. 1E,F**). The number of OROV infected cells at 24 hpi was reduced at least 10-fold in the KO line (**Fig. 1E**).

### The Lrp1-binding chaperone protein RAP inhibits OROV infection of cell lines from taxonomically distinct species

RAP (receptor-associated protein, or Lrpap1) is a high-affinity Lrp1 ligand and critical chaperone of Lrp1 and other LDL receptor family members (Bu, 2001). Domain 3 of RAP specifically binds to two extracellular cluster domains of Lrp1 (CL_II_ and CL_IV_) (**Fig. 2A**) and competes for attachment with other compatible ligands while chaperoning the protein through the ER to the cell surface (Bu, 2001; Ganaie *et al*., 2021). Here, we tested the ability of exogenous mouse RAP_D3_ (mRAP_D3_) to block OROV infection by adding it before infection, at the time of infection, or following infection (**Fig. S2A**). After determining that pre-treatment most effectively blocked infection, mRAP_D3_ was added to murine BV2 microglial cells, monkey Vero E6 kidney cells, and human SH-SY5Y neuroblastoma cells at various concentrations 1 hour prior to infection with 0.1 MOI of OROV, RVFV, or ZIKV as comparators (**Fig. S2B; Fig. 2B-C**). At 24 hpi, samples were tested for infectious virus by plaque assay. As a control, mutant mRAP_D3_ in which two lysines were changed (**Fig. 2A**) was tested in parallel, as these mutations have been shown previously to reduce binding of RAP to CL_II_ and CL_IV_ of Lrp1 (Ganaie *et al*., 2021; Migliorini et al., 2003; Rauch et al., 2020). In all cell lines, treatment with mRAP_D3_ significantly reduced OROV infection compared to untreated cells at all concentrations, and the reduction in infection was comparable to RVFV (**Fig. S2B; Fig. 2B,C**). The mutant mRAP_D3_ was less effective at inhibiting both viruses at lower concentrations, with intermediate levels of inhibition at higher concentrations. ZIKV does not efficiently infect BV2 cells but does infect SH-SY5Y and Vero cells (**Fig. S1A**). Neither mRAP_D3_ nor mutant mRAP_D3_ inhibited ZIKV infection of SH-SY5Y or Vero cells (**Fig. 2B,C**).

**Figure 2.**
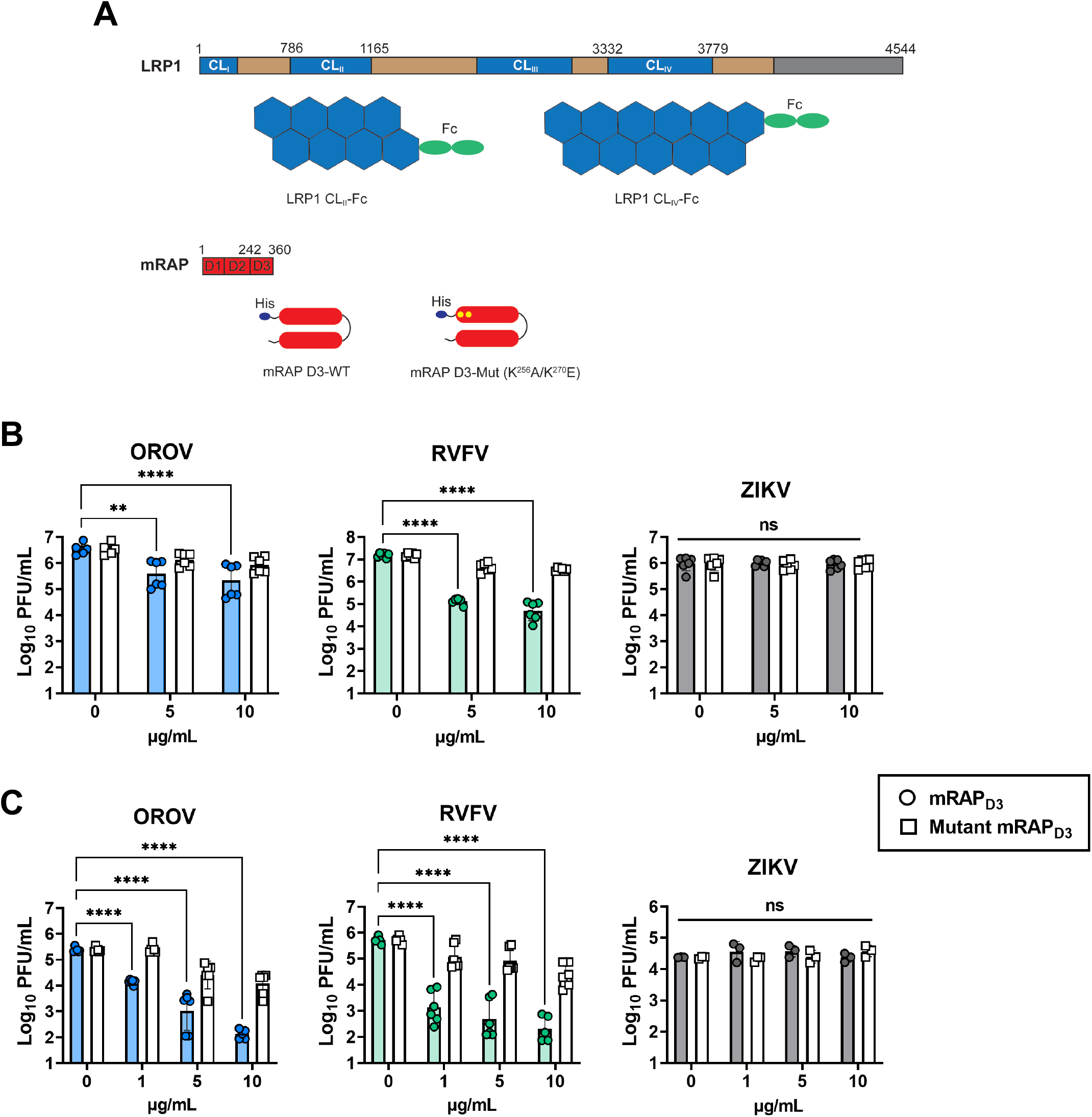
The Lrp1-binding chaperone RAP can inhibit OROV infection of Vero E6 cells and undifferentiated SH-SY5Y cells. (A) LRP1 consists of a 515 kDa extracellular alpha chain (blue/tan) and an 85 kDa intracellular beta chain connected by a transmembrane domain (grey). The alpha chain is further divided into 4 complement-type repeat clusters (CLI-IV; blue), and EGF-like and YWTD domains (tan). Recombinant Fc-fused LRP1 CLII and CLIV were expressed and purified from Expi293 cells for experiments presented here. RAP is a 39 kDa ER-resident consisting of 3 domains (D1-3) that chaparones LDLR family protein including LRP1. Recombinant mRAP D3 WT and mRAP D3 Mut (K256A, K270E) were expressed and purified from BL21 (DE3) cells utilizing an N-terminal His-tag. mRAP_D3_ or mutant mRAP_D3_ was added to (B) Vero E6 nonhuman primate cells, or (C) SH-SY5Y human neuroblastoma cells 1 hour prior to infection with MOI 0.1 of RVFV, OROV, or ZIKV. Samples were harvested at 24 hpi (for RVFV and OROV) or 48 hpi (for ZIKV) and infectious virus was measured by plaque assay or qRT-PCR. Statistical significance was determined using 2-way ANOVA on log-transformed data. Experiments were repeated three times.**, p<0.01; ****, p<0.0001.

### Lrp1 cluster domains CL_II_ and CL_IV_ inhibit OROV infection

Many ligands of Lrp1 bind to the CL_II_ and CL_IV_ extracellular domains, including mRAP_D3_. Recombinant Fc-fusions of the Lrp1 CL domains were shown to block RVFV infection (Ganaie *et al*., 2021). We treated Vero E6 cells with soluble Fc-fused CL_II_ and CL_IV_ proteins (Ganaie *et al*., 2021) and compared the relative infection to untreated cells and Fc-control treated cells. We observed that CL_II_ and CL_IV_ treatment significantly reduced OROV infection compared to the Fc-control treated cells (**Fig. 3A**). These results are comparable to that of treated cells infected with RVFV at the same MOI (**Fig. 3B**). Additionally, ZIKV infection of Vero E6 cells was unaffected by treatment with any Fc-bound Lrp1 proteins (**Fig. 3C**).

**Figure 3.**
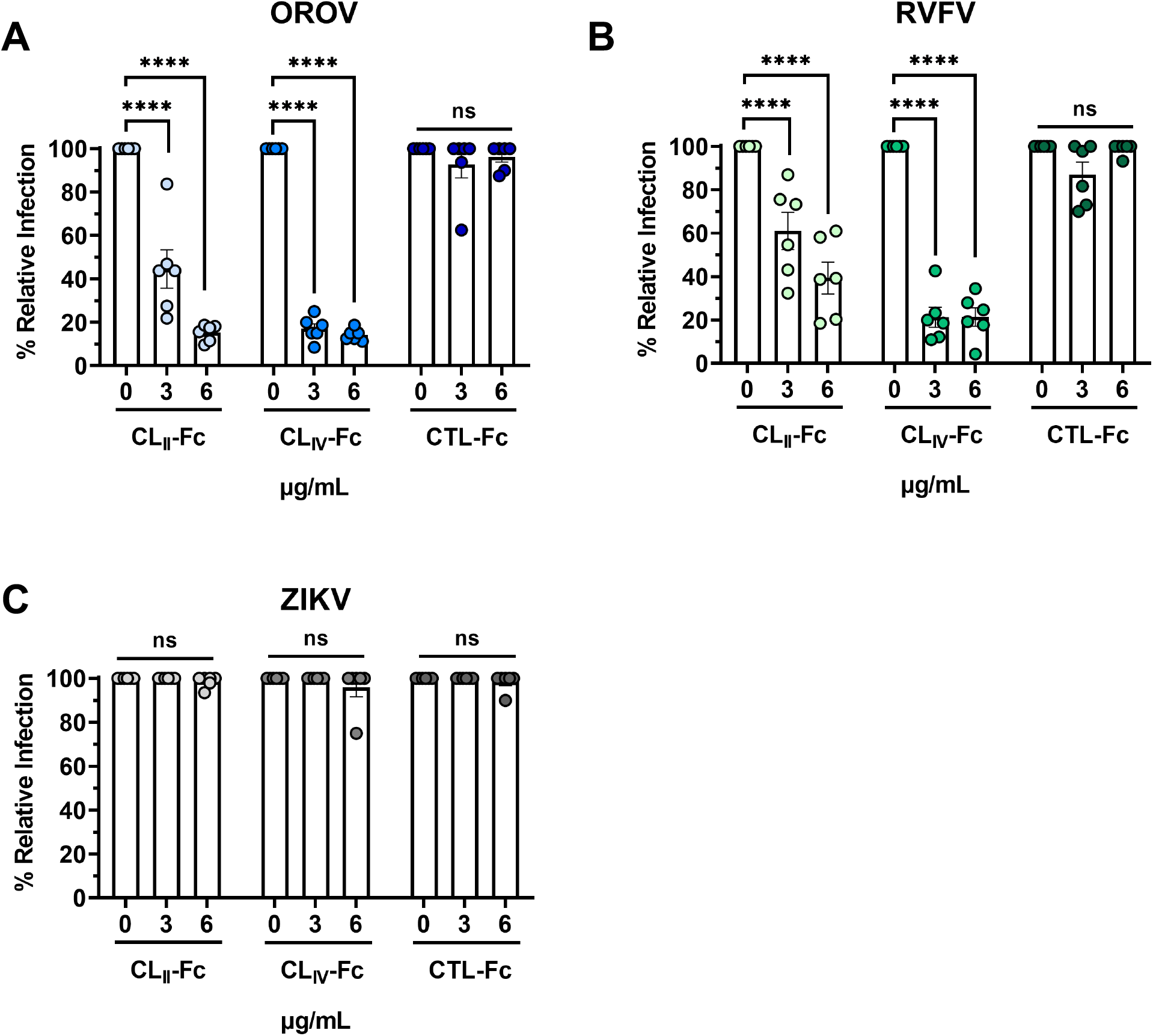
Soluble Fc-bound Lrp1 CLII and CLIV inhibits cellular infection by OROV. (A) Soluble Fc-bound CLII, CLIV, or Fc control proteins were added to Vero E6 cells 1 hour prior to infection with (A) OROV, (B) RVFV, or (C) ZIKV at MOI 0.1. Samples were harvested at 24 hpi (OROV and RVFV) or 48 hpi (ZIKV) and virus was measured by plaque assay and qRT-PCR. Data are expressed as a % of untreated control titers. Statistical significance was determined using 2-way ANOVA. Experiments were repeated two times. ****, p<0.0001.

## Discussion

Here, we show that the conserved protein Lrp1 is a host factor that mediates cellular infection by OROV. The LDLR family and related proteins have recently been implicated in the binding and entry of diverse viruses. LDLR has been reported to play a binding and entry role for Dengue virus (Tree et al., 2019), Hepatitis C virus (Molina et al., 2007), vesicular stomatitis virus (VSV) (Nikolic et al., 2018), and rhinovirus (Hofer et al., 1994). The LDLR-related protein LDLRAD3 mediates entry of Venezuelan equine encephalitis virus (Clark et al., 2021). VLDLR and ApoER2 were recently identified as entry receptors for multiple alphaviruses, including Semliki Forest virus, Eastern Equine encephalitis virus and Sindbis virus, despite differences in E2/E1 amino acid sequence homology (Clark *et al*., 2021). Rift valley fever virus (RVFV) was the first member of *Bunyavirales* found to utilize the LDLR family machinery for entry (Ganaie *et al*., 2021), and evidence suggests Lrp1 may be important for SARS-CoV-2 as well (Devignot et al., 2022). Our studies here reveal that the orthobunyavirus OROV also uses Lrp1 to gain entry. OROV Gn binds similar and potentially overlapping regions within Lrp1 CL_II_ and CL_IV_ domains as phelbovirus RVFV Gn, suggesting some structural similarities despite wide sequence diversity. This study underscores that LDL receptors, such as Lrp1, may play a broader role in cell entry of bunyaviruses than previously appreciated. Studies to characterize the binding and internalization mechanisms of both OROV and RVFV are underway. Whether Lrp1 is used by other members of the *Bunyavirales* order or whether this observation is the result of convergent evolution between OROV and RVFV will be a subject of our ongoing efforts to evaluate the impact of LDL family receptors as viral entry factors and as therapeutic targets.

## Materials and Methods

### Biosafety

Work with OROV and ZIKV was completed in a BSL-2 laboratory following all University biosafety guidelines. Work with RVFV was completed at BSL-3 in the Regional Biocontainment Laboratory (RBL) at the University of Pittsburgh. The authors have approval from the Federal Select Agent Program (FSAP) to work with RVFV in the described facilities.

### Cells

The LPR1^KO^ R4 and RAP^KO^ A7 were generated as previously described (Ganaie *et al*., 2021). All BV2 cells were cultured in Dulbecco’s Modified Eagle Medium (DMEM) (ATCC, 30-2002) supplemented with 1% Pen/Strep, 1% L-Glut, and either 2% (D2), 10% (D10), or 12% (D12) fetal bovine serum. SH-SY5Y and Vero cells were obtained from ATCC (CRL-2266, CRL-1586) and were cultured in D12/F12 media (ATCC, 30-2006) supplemented with 1% Pen/Strep and 1% L-Glut. HEK293T, A549, and N2a clonal KO cells were generated by CRISPR/Cas9 using ribonucleoprotein complexes of Cas9 and *Lrp1*-specific gRNAs, as previously described (Ganaie *et al*., 2021). Resulting cells were sub-cloned and subjected to NGS analysis and STR profiling to confirm the deletion and homogeneity of the clones. HEK293T Lrp1 KO cells were maintained in D10 media, A549 Lrp1 KO cells were maintained in D12/F12 media, and N2a Lrp1 KO cells were maintained in Eagle’s Minimum Essential Medium (ATCC, 30-2003) with 10% FBS.

### Viruses

The BeAn19991 strain of OROV was rescued through reverse genetics and was generously provided by Paul Duprex and Natasha Tilston-Lunel (CVR). RVFV ZH501 was rescued through reverse genetics and provided by Stuart Nichol (CDC). The PRVABC59 (Human/2015/Puerto Rico) strain of ZIKV was obtained from BEI Resources (NR-50240) from the Arbovirus Reference Collection, CDC Fort Collins, CO, USA. VSV-OROV virus was generated as described previously (Stubbs et al., 2021). All viruses were propagated in Vero E6 cells (ATCC, CRL-1586) with standard culture conditions using D2 media supplemented with 1% Pen/Strep and 1% L-Glut. A standard viral plaque assay (VPA) was used to determine the titer of the stocks. The agar overlay for the VPA is comprised of 1x minimal essential medium, 2% FBS, 1% pen/strep, 1% HEPES buffer, and 0.8% SeaKem agarose; the assay incubates at 37C for 4 days (OROV) and 3 days (RVFV) followed by visualization of plaques with 0.1% crystal violet.

### Antibodies

The following antibodies were used: rabbit anti-Lrp1 (abcam ab92544), mouse anti-Lrp1 (Santa Cruz, sc-57353), rabbit anti-OROV N (Custom Genescript), and mouse anti-RVFV N (BEI Resources, NR-43188) for fluorescence immunostaining, and rabbit anti-Lrp1 (Cell signaling, 64099S) and rabbit anti-GAPDH (Thermo Fisher, PA1-987) for western blots.

### Neutralization assays with mRAP_D3_ and Lrp1 clusters

Cells were seeded in a 24-well plate at a density of 2.4E5 cells/mL in D10 the day prior to infection. The following day the media was removed and virus diluted in D2 (MOI 0.1) was added to the cell monolayer in a 200 μL volume. Following a 1-hour incubation at 37 °C, the inoculum was removed. The monolayer was washed with 1X dPBS, and fresh D2 media was added to each well. At the designated timepoint(s) supernatants were collected and infectious virus titers were determined by VPA. In the treatment assays, mRAP_D3_, mutant mRAP_D3_, Fc-CL_II, IV_ or Fc-control (Trastuzumab) were diluted in D2 and added to the cell monolayer followed by a 1 hour incubation at 37 °C. Following the incubation, virus diluted in D2 (MOI 1, 0.1) was added to the media and incubated for 1 hour at 37 °C. After the absorption period, the inoculum was removed, the cell monolayer was washed once with 1x dPBS, and D2 media containing the designated proteins was added in a 500 μL volume. Supernatants were collected at 24 hpi (OROV and RVFV) or 48 hpi (ZIKV) and viral titers were determined through VPA. Vial titers for RVFV and ZIKV were also analyzed through qRT-PCR as previously described (Carossino et al., 2017; McMillen et al., 2018; Waggoner and Pinsky, 2016).

### Immunofluorescence

Cover glass (FisherBrand, 12-546-P) was sterilized in 70% EtOH and coated with BME Cultrex (R&D Systems, 3432-010-01) prior to seeding. Cells were seeded the day prior to staining at a density of 1E5 cells/mL in D10 media. Upon harvest, virus-infected cells were fixed in 4% PFA for 20 minutes, followed by permeabilization with 0.1% Triton X-100 detergent + 1xPBS for 15 minutes at room temperature (RT). Cells were blocked using 5% normal goat serum (Sigma, G9023) for 1 hour at RT, followed by incubation with the primary antibodies rabbit anti-LRP1 at 1:200 (abcam, ab92544), 1:50 mouse anti-Lrp1 (Santa Cruz, sc-57353) mouse anti-RVFV N 1:200 (BEI, NR-43188), or rabbit anti-OROV N 1:200 (Genescript Custom) for 1 hour at RT. The secondary antibodies goat anti-mouse Cy3 (JacksonImmuno, 115-165-003), goat anti-rabbit 488 (JacksonImmuno, 111-545-003), goat anti-mouse 488 (JacksonImmuno, 115-545-003), or goat anti-rabbit Cy3 (JacksonImmuno, 111-165-144) were added (1:500 dilution) for 1 hour at RT. The cells were counterstained with Hoescht and mounted using Gelvatol.

### Protein expression and purification

Recombinant protein expression and purification was performed as described previously (Ganaie 2021). Briefly, expression plasmids containing mRAP_D3_ or mutant mRAP_D3_ were transformed in BL21(DE3) E. coli cells (Novagen), cultured in Luria Broth media at 37 °C to an OD600 of 0.6 and induced with 0.5 mM isopropyl-b-D-thiogalactoside (IPTG) for 12 hr at 18 °C. Cells were harvested, resuspended in lysis buffer containing, and lysed using an EmulsiFlex-C5 homogenizer (Avestin). The resulting pellet was resuspended in buffer containing 2 M urea, 20 mM Tris-HCl (pH 8.0), 500mM NaCl, 2% Triton X-100 prior to centrifugation at 47,000 x *g* at 4 °C for 10 mins. Inclusion bodies were isolated after repeated rounds of resuspension in urea and centrifugation. The final pellet was resuspended in 20 mM Tris-HCl (pH 8.0), 500 mM NaCl, 5 mM imidazole, 8 M urea, and 1 mM 2-mercaptoethanol and refolded on a NiFF (GE Healthcare) column using a reverse linear urea gradient with imidazole. Endochrome-K kit (Charles River) was used, following the manufacturer’s instructions to determine endotoxin levels for purified mRAP_D3_ proteins and the control protein.

Expression plasmids encoding CL_II_ or CL_IV_ with an Fc- and His_6_-tag were transfected in Expi293F expression cells (Gibco-Thermo Fischer Scientific) using the ExpiFectamine293 transfection kit. Cells were cultured in Expi293 Expression Media at 37 °C and 8% CO_2_ for 5 days. Cells were separated from supernatant by centrifugation and supernatant containing the secreted proteins was loaded onto a NiFF (GE Healthcare) column as described above.

## Acknowledgements

We thank members of our research groups for helpful discussions and support. We thank W. Paul Duprex and Natasha Tilston-Lunel (University of Pittsburgh) for providing the OROV virus used in these studies. The authors also thank Stacey Barrick for coordinating housing for animal studies. The following reagents were obtained through BEI Resources, NIAID, NIH: Zika Virus, PRVABC59, NR-50240; RVFV anti-N mAb, Clone 1D8, NR-43188. Research was supported by NIH grants (R01AI161765 and R21AI163603 to G.K.A and A.L.H; R01NS101100 to A.L.H; P01AI120943 to G.K.A.; R01AI107056 to D.W.L.; R01AI130152 to T.E.)

## Author Contributions

Investigation and Validation, all authors; Methodology, all authors; Writing – initial draft M.M.S and A.L.H.; Writing – review and editing, G.K.A, D.W.L.; Supervision, A.L.H, D. W. L., G.K.A.; Funding Acquisition, A.L.H, D. W. L., G.K.A.

## Supplemental Figure Legends

**Supp. Fig 1.**
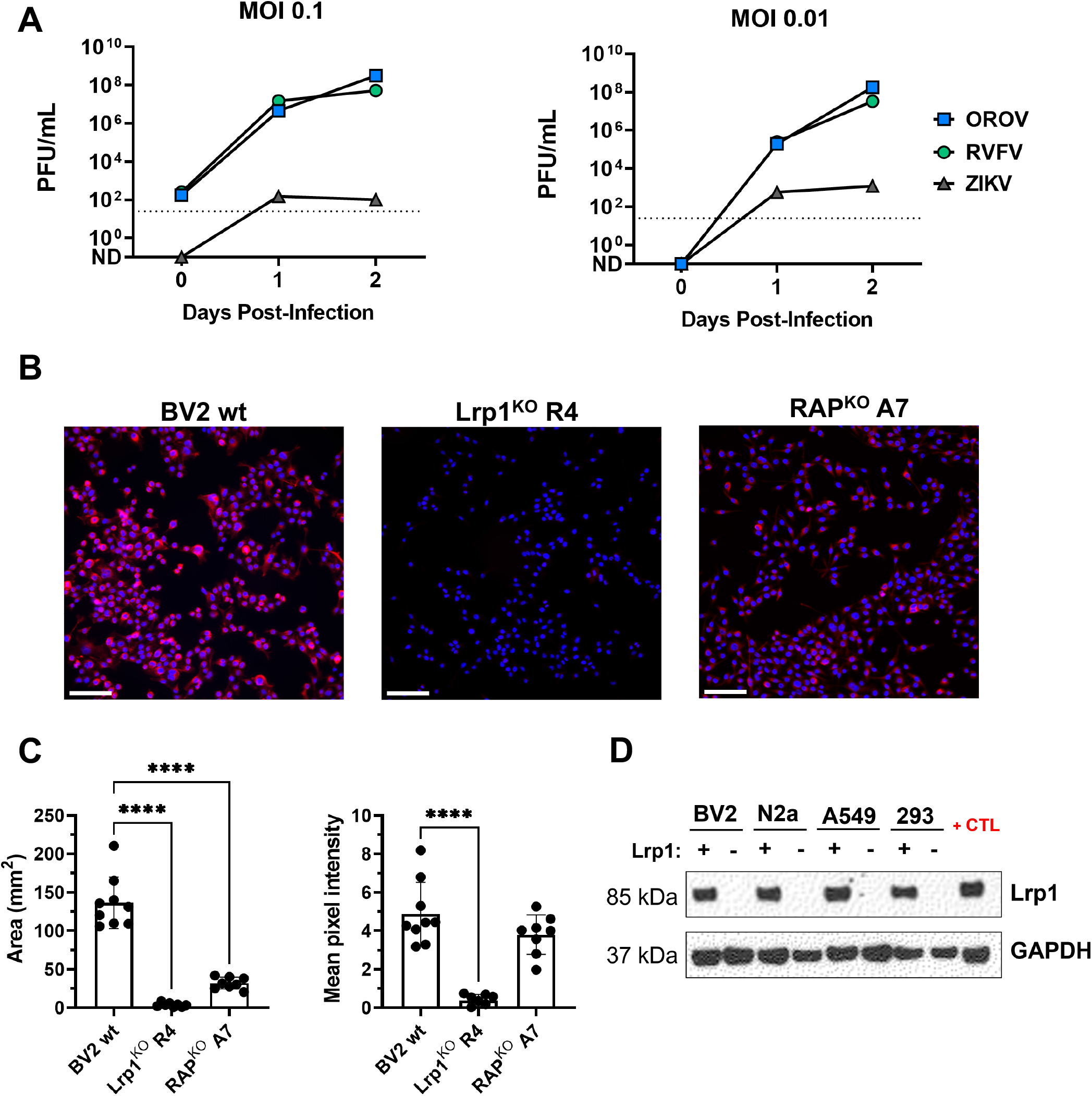
OROV, RVFV, and ZIKV replication in BV2 cells and Lrp1KO cell lines. (A) Growth of OROV, RVFV and ZIKV in wild-type mouse microglial BV2 cells at MOI 0.1 and MOI 0.01. Viral titers determined by plaque assay. (B) BV2 Lrp1KO and RAPKO cell lines described in Ganaie et al. 2021 show reduced expression of Lrp1 based on immunofluorescence with anti-Lrp1 mAb (20X). Scale is 50μm. (C) Image quantification of cells from panel (B) shows significantly less area (mm2) and mean pixel intensity associated with Lrp1 staining in KO cell lines. (D) Western blot for Lrp1 demonstrating KO in BV2, N2a, A549, and HEK293T (293) cell lines. Positive control (+CTL) is homogenized C57BL/6J mouse liver. Statistical significance was determined using 2-way ANOVA. ****, p<0.0001.

**Supp. Fig. 2.**
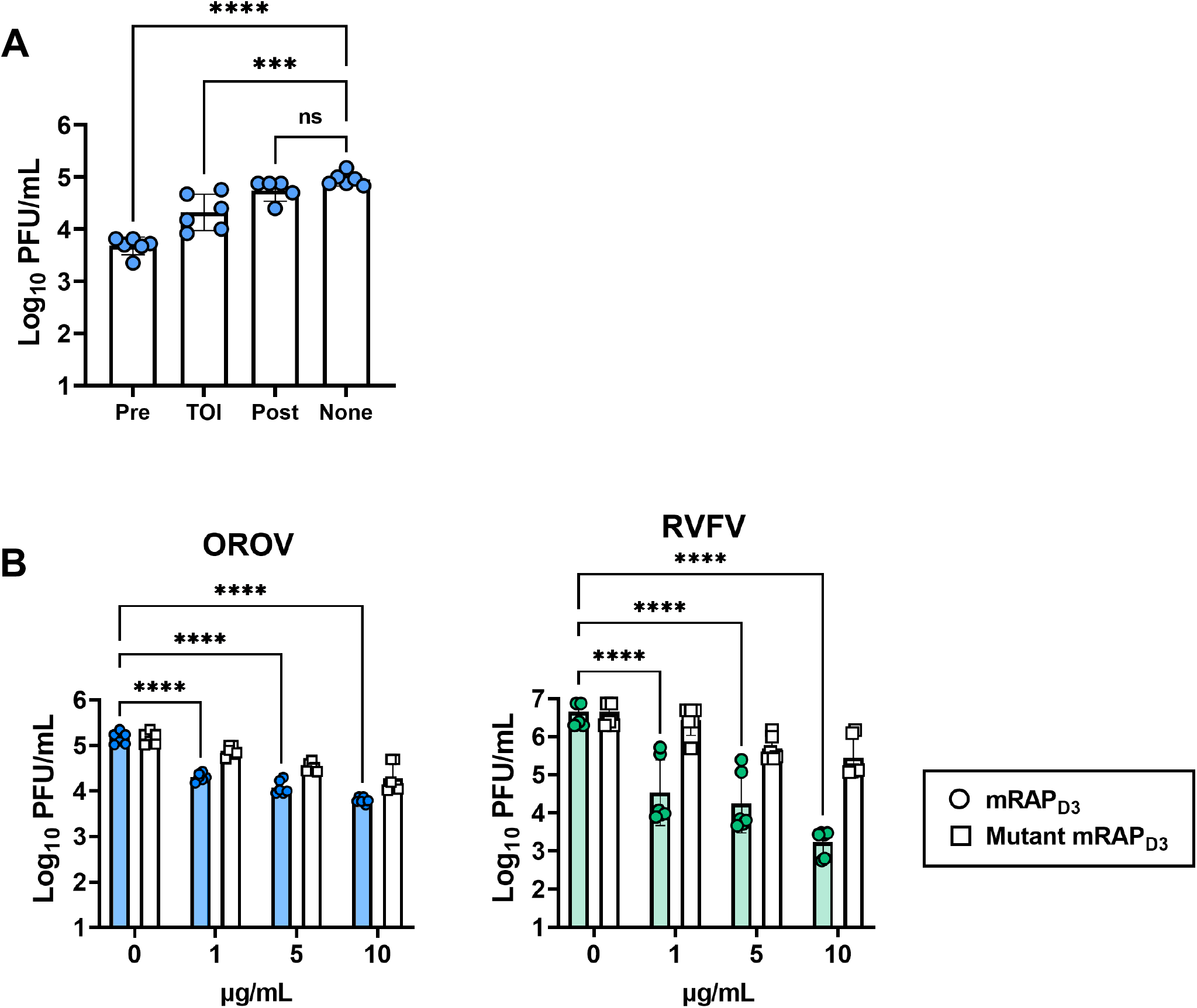
The Lrp1-binding chaperone RAP can inhibit OROV infection of Vero E6 cells and BV2 mouse microglia cells. (A) 10μg/mL mouse RAP D3 (mRAP_D3_) was added to Vero E6 cells 1 hour prior to infection (Pre), at the time of infection (TOI), or after the infection (Post) with MOI 0.1 of OROV. Samples were harvested at 24 hpi and infectious virus was measured by viral plaque assay. (B) mRAP_D3_ or mutant mRAP_D3_ were added to BV2 cells 1 hour prior to infection with MOI 0.1 of OROV or RVFV. Samples were harvested at 24 hpi and infectious virus was measured by plaque assay. Statistical significance was determined using 1-way ANOVA or 2-way ANOVA on log-transformed data. Experiments were repeated two times. ***, p<0.001; ****, p<0.0001.

